# A Cross-Platform Comparison of Student’s Perceptions of the Learning Environment in an Introductory Microbiology Course

**DOI:** 10.1101/274571

**Authors:** Melissa V. Ramirez, Virginia S. Lee, Lindsay Hamm, Michael E. Taveirne, Alice M. Lee

## Abstract

**Abstract:** Students’ positive perceptions of the learning environment increase retention and persistence in STEM disciplines. This study presents results from a Learning Environment Questionnaire administered at the beginning and end of the semester in a redesigned general/introductory microbiology course offered in traditional lecture, flipped and online sections. Split-sample t-tests and chi-squared tests were used for cross-section and within section analyses, respectively. The findings support the study hypothesis that student perceptions of the learning environment will vary as a function of platform. This work demonstrates the additional effect of the time in the semester when students complete the questionnaire and this effect on students’ perceptions over the course of the semester relative to their initial perceptions. The results of this study offer insight on student perceptions of the learning environment as universities embrace online education in introductory and gateway courses as a response to rising student enrollments and diminishing resources.

## INTRODUCTION

Across the country, campuses are redesigning STEM gateway courses to improve student performance, retention, time to graduation, and to encourage continued study in scientific disciplines. To accommodate both student preferences and swelling university enrollments, often these courses are offered in three platforms: traditional lecture, blended (or flipped), and fully online. Because they offer a cost effective way to teach more students and enable non-traditional students and distant students to enroll, online courses, in particular, have received a great deal of scrutiny recently.

While some researchers have noted the limitations of cross-platform comparisons (1, 2, 3), there is a vigorous strand of research that compares students’ performance and perception of the learning environment across traditional, flipped and/or online platforms (see, for example, the U.S. Department of Education, Office of Planning, Evaluation and Policy Development, meta-analysis of ninety-nine cross-platform studies (4)). As campuses and course providers such as Coursera and EdX have accumulated more experience with student behavior and performance in online courses, they have found that typically more mature learners who are self-directed persist and perform better in online classes than traditional-aged college students (5, 6). More recently, as researchers have come to appreciate the links between emotion and cognition (7, 8), educators have paid increasing attention to students’ perception and experience of the classroom as important indicators of satisfaction, a sense of belonging, retention beyond the first year, and persistence in the discipline (9, 10). Educators and researchers have used various ways of measuring student learning (11, 12, 13, 14, 15, 16, 17), some more robust than others, as well as students’ perception of the learning environment (11, 13, 15, 16, 17).

The study described in this article assesses students’ perceptions of the learning environment of a redesigned general microbiology course offered in three platforms.

### Project Description

General Microbiology (MB 351) is a large enrollment (greater than 1,000 students/year) introductory course taught year-round in multiple sections by different instructors through the Department of Biological Sciences at North Carolina State University. The course is required for Microbiology majors as well as other life science disciplines. Originally, the course was offered as a 250-student traditional lecture course taught by one instructor in an auditorium-style classroom with PowerPoint lecture slides.

Following a substantial redesign of the course, MB 351 is currently offered in three modes: a smaller traditional lecture with approximately 50-75 students, flipped (60-90 students), and fully online (approximately 250 students). Flipped sections are taught in a specially designed Student Centered Active Learning Environment using Upside-down Pedagogies (SCALE-UP) classroom (18). Extensive use of in-class activities and active learning techniques (e.g., clickers, case studies, think-pair-share, problem solving) are employed in the flipped class. The online section utilizes weekly assigned mini-lectures (about 15-20 minutes in length, 4 times per week), supporting animations, case studies, discussion forums, and a number of summative assessments that mirror those in the flipped section. All sections utilize online resources to some degree.

A multi-methods assessment of the newly redesigned course was conducted in spring 2016. The assessment project comprises a pre-/post-direct measure of student learning, a pre-/post -survey of students’ perception of their MB 351 section as a learning environment, student focus groups, and instructor interviews. Following submission of an application, the study was exempted from IRB review. The goals of the study were to illuminate the dynamics of the newly designed course, including student performance and student perception of the learning environment, to compare differences across the three platforms, and to inform further revision of the course by the instructors.

### Hypothesis

This article focuses on students’ perception of the course as a learning environment across the three learning platforms. The associated study hypothesis is that students’ perceptions of the learning environment will vary as a function of the platform.

## METHODS

### Development of the Survey Instrument

Insights gained from student focus groups conducted during a spring 2014 pilot assessment project led to a desire to examine students’ perceptions more rigorously through the development and administration of a survey instrument in spring 2016.

As part of the survey development, a selective review of the research literature on classroom environment questionnaires, intensive examination of an instrument developed by colleagues at Purdue University (Levesque- Bristol, C. and Nelson, D., unpublished data), and adaptation of an instrument developed as part of a study of large classes at UNC-Chapel Hill (Lee, V.S. and Williford, L.E., unpublished data) was performed.

The resulting Learning Environment Questionnaire (LEQ) is a dual-response, online survey containing four sub-scales administered early (Pre-LEQ) and late (Post-LEQ) in the spring 2016 semester through Qualtrics. The four sub-scales—Academic Environment, Affective Dimension, Self-Determination, and Perceived Value of the Material—measure different dimensions of the classroom environment and experience that may affect student performance, motivation, and persistence in the discipline. Rather than creating our own instrument *de novo*, we adapted scales whose reliability and validity had already been confirmed. The Academic Environment and Affective Dimension sub-scales from comparable scales developed for the UNC-Chapel Hill large class study and the Self-Determination and Perceived Value of the Material sub-scales from the Purdue University instrument (see Appendix 1 for the Learning Environment Questionnaire including the associated statements for each sub-scale).

Students rated each item twice, each time on a four-point Likert scale according to its Importance for their Learning and the Degree to Which It Described Their Class. Consequently, each item yielded an Importance (I) rating, a Class (C) rating and a Class minus Importance (C – I) rating.

### Sampling

We emailed the questionnaire to 274 students registered in MB351 at the beginning of the semester, of whom 157 completed it for a response rate of 57%, and 271 students at the end of the semester, of whom 115 completed it for a response rate of 42%. The lower number of students mailed at the end of the semester was due to attrition.

### Analysis

Following the administration of the questionnaire, we imported the questionnaire data into STATA for analysis. We removed eighteen cases from the Pre-LEQ and sixteen cases from the Post-LEQ, because these students only provided the section in which they were enrolled and then answered no other questions.

We left the remaining cases in the dataset, even those with some missing data due to unanswered questions, in order to use as much original data in the analysis as possible. Then we list wise deleted cases with missing data from the sample in t-tests only for the questions students did not answer. We calculated the means of the Class and Importance ratings for each item in each of the four sub-scales (“rowmean” in STATA). Then we calculated the Class - Importance ratings by taking the mean of the values obtained by subtracting each case’s Importance rating from its corresponding Class rating for each item comprising the subscale.

Participation in both the Pre- and Post-LEQ was voluntary, and students were only asked to indicate in which course section they were enrolled, but not any further identifying information. As a result, we cannot make direct comparisons between students across the Pre- and Post-LEQs. Instead, we used split-sample t-tests in our cross-section analyses and chi-squared tests to compare data within the sections. We also employed Fisher’s Exact tests when observed frequencies were lower than five. Again, we used a p value of .05 to determine significance.

## RESULTS

Tables 1 - 3 below summarize the results of the Pre- and Post-LEQ survey. Table 1 summarizes the between-section differences on the Pre- and Post-LEQ ratings. Table 2 provides the within-section comparisons on the Pre- and Post-LEQ ratings. Table 3 isolates the significant Pre- and Post-LEQ class-minus-importance ratings within sections from Table 2 for the sake of clarity. Table 4 presents the total number of statistically significant differences by section across the three tables.

**Table 1.** Significant Differences on Pre- and Post-LEQ Sub-Scale Ratings Between Sections.

T = Traditional Lecture, F = Flipped, O = Online
|  | **T vs O** |  | **T vs F** |  | **O vs F** |  |
| --- | --- | --- | --- | --- | --- | --- |
|  | **Pre-** | **Post-** | **Pre-** | **Post-** | **Pre-** | **Post-** |
| **Academic Environment** |  |  |  |  |  |  |
| Importance | -.08 | .03 | -.02 | -.05 | .06 | -.08 |
| Class | -.17 | .07 | .10 | .20 | .27* | .13 |
| Class – Importance | -.09 | .08 | .12 | .23 | .21 | .15 |
| **Affective Dimension** |  |  |  |  |  |  |
| Importance | .20 | .36* | .08 | -.09 | -.12 | -.45* |
| Class | .0 | .2 | -.12 | -.25 | -.12 | -.45* |
| Class – Importance | -.19 | -.13 | -.19 | -.18 | .0 | -.05 |
| **Self Determination** |  |  |  |  |  |  |
| Importance | -.03 | .16 | .07 | -.15 | -.04 | -.31* |
| Class | .88 | .35* | .09 | -.02 | -.79 | -.37* |
| Class – Importance | .91 | .12 | .16 | .12 | -.75 | .1 |
| **Perceived Value** |  |  |  |  |  |  |
| Importance | .05 | .04 | .10 | -.22 | .05 | -.26* |
| Class | .88 | -.01 | .09 | -.24 | -.79 | -.23 |
| Class - Importance | .83 | .01 | -.01 | -.01 | -.84 | -.02 |

\*Indicates significant difference

**Table 2.**
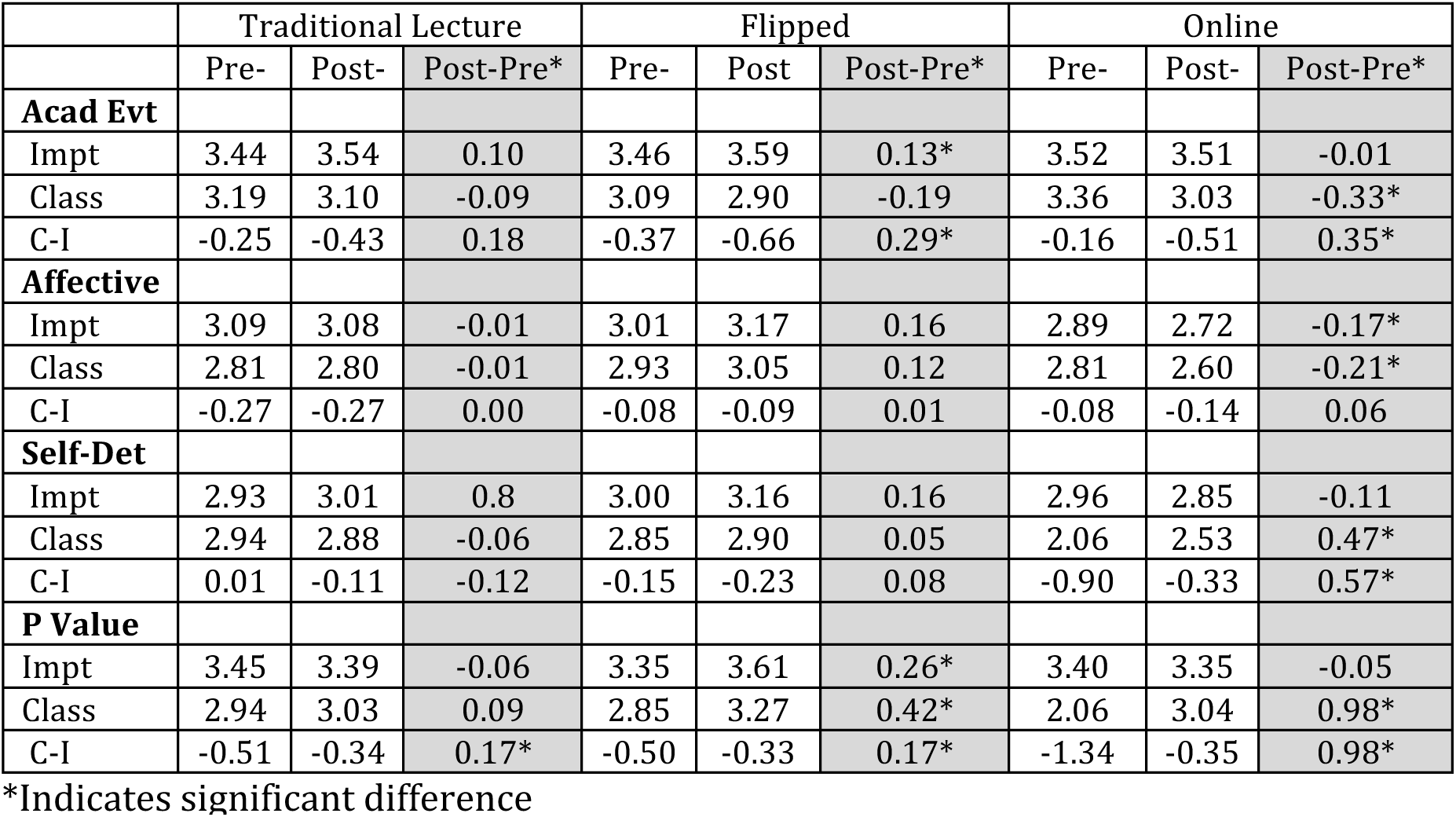
Comparison of Pre- and Post-LEQ Ratings within Sections.

**Table 3.** Significant Pre- and Post-LEQ Class-minus-Importance Sub-Scale Rating Differences within Sections.

|  | Traditional |  | Flipped |  | Online |  |
| --- | --- | --- | --- | --- | --- | --- |
|  | Pre- | Post- | Pre- | Post- | Pre- | Post- |
| **Academic Environment** | (.25)* | (.43)* | (.37)* | (.66)* | (.16)* | (.51)* |
| **Affective Dimension** | (.27)* | (.27)* | ns | ns | ns | ns |
| **Self Determination** | ns | ns | ns | ns | (.90)* | (.33)* |
| **Perceived Value** | (.51)* | (.34)* | (.50)* | (.33)* | (1.34) | (.35)* |

\*Indicates significant difference

**Table 4.**
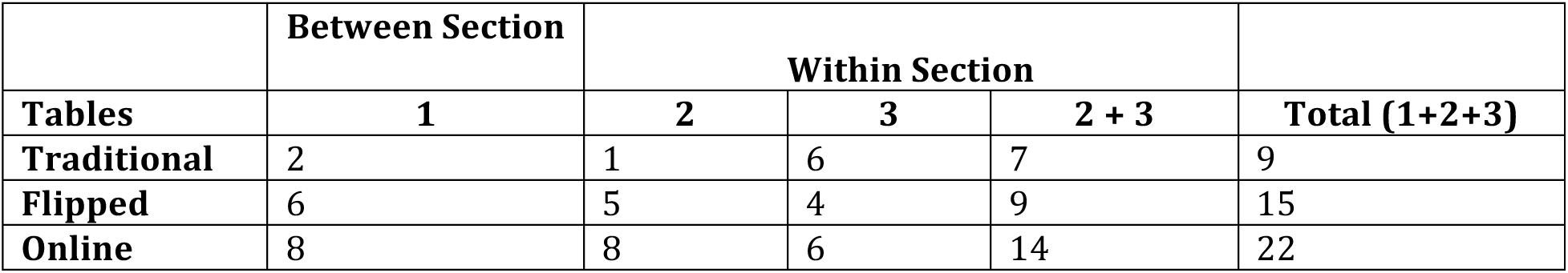
Summary of Unique Statistically Significant LEQ Effects by Section.

Administration of the LEQ early in the semester allowed us to capture students’ initial impressions of the classroom environment. In contrast, administration of the LEQ again at the end of the semester allowed us to capture students’ final impressions of the classroom environment, as a result of their actual experience of the course during the semester. Differences between Pre- and Post- ratings are important, because the differences capture the effect of students’ experience of the course relative to their initial expectations and first impressions.

In addition, because students rated each statement on the survey according to both its importance to their learning (I) and the extent to which it occurred in their section (C), we were able to capture not only students’ perception of the class itself, but also how their experiences over the course of the semester shaped their sense of what was important to their learning. A negative C – I rating indicates that the classroom is lacking relative to students’ judgment of the item’s importance for their learning, and a positive C – I rating that the item exists in the classroom beyond students’ judgment of its importance for their learning.

### Comparisons between Sections on the Pre- and Post-LEQ

Comparisons among sections are complex. Between any two sections, differences could reflect differences in the students in the sections, the instructors themselves including their personalities, instructor relationship with students, classroom practices, teaching experience, the nature of the learning platform, and students’ predisposition to favor one platform over another. In their classic articles, Ross and Morrison (3, 19) called for a “happy medium” in educational technology research by balancing high rigor of studies (or “internal validity”) with relevance to real world applications (or “external validity”). The current study attempts to strike that balance.

For these reasons, a single inter-sectional statistically significant effect may be less noteworthy than a consistent <u>pattern</u> of statistical effects between and among sections. The following section describes two such a patterns from the study results and offers an explanation of the patterns.

### Between Section Differences on the Pre-LEQ

There are good reasons to assume that students taking the course in different platforms would have different perceptions and expectations about their class as a learning environment even early on in the semester. In terms of the four sub-scales of the LEQ, on Academic Environment, many students’ negative perceptions of online learning (20, 10) could result in lower Class ratings related to the instructor’s ability to communicate expectations clearly, develop exams and assignments that let students demonstrate what they learn, and so on. Certainly we would expect differences in the Affective Dimension—including instructor’s knowledge of who students are, that they are trying, that students will have a chance to participate in class, get to know other students, and so on— between the online section and both the traditional and flipped conditions in which instructor and students regularly meet face-to-face. On Self Determination, while all platforms could offer students choices and options, conveying confidence in students’ ability to do well, listening to how students would like to do things, and trying to understand how students see things should be far more difficult in the online condition; there is no regular face-to-face contact and instructional design decisions made during the development of the course are harder to change while the semester is in progress than they are in face-to-face environments. Similarly, in terms of Perceived Value, instructors of online courses could find it more challenging to convey the value of the material learned and its relationship to future study and careers than in face-to-face settings.

Further, we might expect particularly strong contrasts between the Traditional and Online sections or the Online and Flipped conditions. Although not formally designed this way, the traditional lecture section functions as a control in this study, representing the customary way of teaching in universities. The Flipped and Online sections on the other hand represent experimental conditions that depart from customary practice. In theory, the flipped condition facilitates deeper forms of student engagement during face-to-face sessions through advanced preparation, all enabled by technology. In addition, the online condition replaces most face-to-face instruction and instructor/student and student/student interaction with virtual instruction and interaction enabled through technology.

In light of all of the above, it is then very surprising that with one exception—the Class rating on the Academic Environment sub-scale between the Online versus Flipped sections, favoring the Online section, there are <u>no</u> significant differences among students’ Class, Importance, and Class minus Importance ratings on any of the sub-scales on the Pre-LEQ (see Table 1).

One possibility is that after the first year, undergraduate students in similar programs would have had comparable educational experiences, the overwhelming majority in traditional lecture courses, and they would carry the assumptions from these experiences into future courses regardless of platform. In the absence of an experience to the contrary, most undergraduate students come to a course with stable assumptions about the importance of certain factors in their learning and the likelihood that they will be present in the current class based on their past experience. In the beginning at least, these prior experiences trump other factors that lead students to make the judgments they do.

Further, in answering the survey questionnaire, many students who complete the survey in the first place may do so hastily. As they complete the questionnaire, they may not be particularly reflective or even thinking about the nuances of teaching and learning in different platforms. Even more so when they complete the survey outside of class online as in the current study.

As we stated above, in addition to the potential effects of platform, there are also student and instructor effects that could balance the anticipated effects of platform, even early in the semester. For example, all of the instructors in the study are non-tenure track Teaching Assistant Professors who are very dedicated to their teaching, spend a lot of time planning and engaging with students, and attending faculty development workshops offered at the University. The instructor of the online section, however, has significantly more teaching experience than the other two instructors: 20 years versus 4 years each for the other two instructors. In previous semesters, whether she was teaching in the traditional or flipped platforms, students have evaluated her highly. As the online students completed the Pre-LEQ, the instructor’s reputation could have balanced out any negative perceptions of online classes.

### Between Section Differences on the Post-LEQ

Differences between the sections do emerge in the Post-LEQ because of students’ actual experience of the course during the semester (see Table 1). All of the significant differences are between the online section and one of the other two sections, and all are in favor of the non-online sections: two differences with the traditional section, and five differences with the flipped section. In addition, the only Pre-LEQ difference on the Academic Environment Class rating between the online and flipped section (and the only between-section difference, pre- or post-, that favored the online section) disappeared in the Post-LEQ. Finally, there are no between-section significant differences between the Traditional and Flipped sections.

### Comparisons within Sections on the Pre- and Post-LEQ

Looking at the within-section differences sheds some light on the between-section differences. As we see in Table 2, students’ Importance and Class ratings remained very stable in the Traditional Lecture section. By contrast, more significant differences emerged in the online section and fewer in the Flipped section. In light of the discussion above, this is not surprising if we assume that the assumptions students brought into the course at the outset of the semester were based largely on their prior course experiences, in which traditional lecture courses predominate. For students in the Traditional section, these assumptions were not challenged and altered by their actual experience of the course in the same way that they were in the online section and, to a lesser degree, the flipped section as “experimental” sections.

Some of the changes were positive. Students’ perception of Perceived Value of the Material showed a positive change over the course of the semester with a very dramatic change in the Online section, although in all three sections a negative discrepancy still existed between students’ Class and Importance ratings. Additionally, in the online section, students’ average Self Determination Class rating increased, leading to a positive change in the Class – Importance rating. Again, this may reflect the expertise of the very experienced instructor of the online section that mitigated the challenges of the platform itself.

It is noteworthy, however, that the online section was the only section in which significant negative pre/post differences occurred on any sub-scale: a decrease in the Class rating on Academic Environment, and decreases on both the Importance and Class ratings on Affective Dimension. Again, however skilled the instructor, over the course of the semester, students experienced the actual challenges of learning in the online environment including its anonymity and impersonality, which they in turn communicated in lower ratings on the associated sub- scales.

Further reflecting the impact of students’ actual experience of the course relative to their entering assumptions is the greater number of significant within-section Pre- and Post-LEQ Class minus Importance ratings differences in the Online section compared to the other two sections (see Table 2). In the Online section, there were significant differences on three of the four sub-scales versus only two for the Flipped and one for the Traditional sections. In addition, as noted above, there were significant negative differences on both the Importance and Class ratings on the remaining sub-scale, Affective Dimension, for the Online section. Since both ratings declined, there was not a corresponding decrease in the Class – Importance rating.

## DISCUSSION

The study findings support the hypothesis that student perceptions of the learning environment will vary as a function of platform (that is, Traditional, Flipped or Online). The findings extend the hypothesis, however, to the extent that they demonstrate the additional effect of the time in the semester at which students complete the questionnaire and the special characteristics of the online section versus the two face-to-face sections and their effect on students’ perceptions over the course of the semester relative to their initial perceptions. They also suggest that individual instructors can mitigate the potentially negative effects of platform on student perceptions to some degree.

As noted above, statistically significant effects in Class-minus-Importance ratings within sections are an indication of either student discontent (negative difference) or disequilibrium between instructor and student assumptions about teaching and learning (positive difference) or both. Very small discrepancies between Class and Importance ratings signal student contentment, perhaps a good thing, but perhaps also an indicator, depending on the sub-scale and particular statement in question, that instructors are not sufficiently challenging students’ expectations. On the other hand, very large negative discrepancies between Class and Importance ratings may indicate a potential problem that bears further investigation by the instructor. The investigation might lead to possible modifications of classroom practices and/or addressing perceptions directly with students. In contrast, large positive discrepancies between Class and Importance ratings may indicate disparities between students’ assumptions about teaching and learning and the instructor’s own assumptions about what is important for their learning.

Further, changes in the size of discrepancies over the course of the semester are also significant: both widening negative and positive discrepancies are red flags that instructors should reflect upon seriously, since they indicate that students’ actual experience of the course has violated in some way their assumptions and expectations. On the other hand, the narrowing of both negative and positive discrepancies indicates a gradual alignment of instructor and student expectations over the course of the semester. In either case, discrepancies provide fertile ground for learning, whether on the part of instructors, students or both.

The study findings suggest that prior classroom experiences, dominated by the traditional lecture in scientific disciplines, shape students’ entering expectations and perceptions of current learning environments. Learning environments that challenge or diverge from these expectations and perceptions may create a sense of disequilibrium and discontent among students as the semester unfolds and the reality of the classroom experience makes itself apparent. Further, even if performance is nearly equivalent across different learning platforms, the learning environments themselves are not necessarily equivalent in terms of the types of classroom experiences they afford and students’ perceptions of those experiences. Experiences that cause disequilibrium and discontent may affect students’ motivation to persist in the course and the discipline and influence their overall level of satisfaction with the university. That recognition was an early stimulus for undergraduate education reform in the sciences including the pioneering work of Sheila Tobias (23).

As the learning environment most radically different from the traditional classroom experience, the online course challenges students’ expectations and perceptions most radically. At the same time, online courses represent the new “large class,” an earlier answer to educating more students with equal or fewer resources, particularly in large public universities. In comparison to large, face-to-face classes, however, the charge for teaching and learning in online environments is even more daunting. The challenges of the large class—anonymity, depersonalization, lack of accountability—are only exacerbated in the online classroom, and best practice in large class teaching— interactive lecture, clickers, small groups, advanced preparation before lecture—is harder to replicate.

The results of this study offer a word of caution as universities embrace online education in introductory and gateway courses such as MB 351 as a response to rising student enrollments and diminishing resources. Since the online environment favors students who are highly self-directed, instructors should develop ways of helping all students succeed in online courses. Creating or using an online learning readiness instrument such as “TOOL” (10) to alert students to the challenges of and prepare them for taking an online course. Provide a weekly student preparation sheet and study guide that focuses on weekly goals and key concepts to master (Students in all sections would benefit from this practice); and offer face-to-face review sessions, study opportunities, and provide an online equivalent such as Blackboard Collaborate sessions for those who cannot attend in person.

In addition, instructors should try to bridge the perspective-divide between students and their entering expectations of teaching and learning with the realities of the online environment. Instructors should explain in the first class session how the course is structured and why it is structured that way, what students can expect over the course of the semester, and what learning requires of them in the course.

Finally, departments should assign instructors to online courses carefully, favoring those who are motivated by the challenges of teaching in the online environment and open to exploration of evolving best practice in the platform.

## Acknowledgements

We would like to thank at North Carolina State University, Distance Education and Learning Technology Applications (DELTA) unit for providing their professional expertise in course redesign; and Robert Beichner, Director of the STEM Education Initiative, for financial support of the course redesign assessment project.

## Supplemental Materials

**Appendix 1:** Learning Environment Questionnaire (LEQ) including the associated statements for each sub-scale

## Notes

**Conflict of Interest Notification Page** Our study funders are internal North Carolina State University sources, specifically the STEM Education Initiative. This initiative provided funding for Virginia S. Lee’s education consultant fees for assessment of student learning in a redesigned General/Introductory Microbiology course (MB351). The supporting source had no involvement in the study design, collection, analysis, and interpretation of the data, in the writing of the report, and in the decision to submit the report for publication.

